# Attention is required for knowledge-based sequential grouping of syllables into words

**DOI:** 10.1101/135053

**Authors:** Nai Ding, Xunyi Pan, Cheng Luo, Naifei Su, Wen Zhang, Jianfeng Zhang

**Affiliations:** College of Biomedical Engineering and Instrument Sciences, Zhejiang Univ., China; Key Labratory for Biomedical Engineering of Ministry of Education, Zhejiang Univ., China; State Key Laboratory of Industrial Control Technology, Zhejiang Univ., China; Interdisciplinary Center for Social Sciences, Zhejiang Univ., China; Neuro and Behavior EconLab, Zhejiang Univ. of Finance and Economics, China; School of International Studies, Zhejiang Univ., China; Mental Health Center, School of Medicine, Zhejiang Univ., China

## Abstract

How the brain sequentially groups sensory events into temporal chunks and how this process is modulated by attention are fundamental questions in cognitive neuroscience. Sequential grouping includes bottom-up primitive grouping and top-down knowledge-based grouping. In speech perception, grouping acoustic features into syllables can rely on bottom-up acoustic continuity cues but grouping syllables into words critically relies on the listener’s lexical knowledge. This study investigates whether top-down attention is required to apply lexical knowledge to group syllables into words, by concurrently monitoring neural entrainment to syllables and words using electroencephalography (EEG). When attention is directed to a competing speech stream or cross-modally to a silent movie, neural entrainment to syllables is weakened but neural entrainment to words largely diminishes. These results strongly suggest that knowledge-based grouping of syllables into words requires top-down attention and is a bottleneck for the neural processing of unattended speech.

## Introduction

Sequentially grouping events into temporal chunks is a fundamental function of the brain (Lashley, 1951, Gavornik and Bear, 2014). During speech comprehension, for example, sequential grouping occurs hierarchically, with syllables being grouped into words and words being grouped into phrases, sentences, and discourses. Similarly, during music perception, musical notes are hierarchically grouped into meters and phrases. Neurophysiological studies show that slow changes in neural activity can follow the time course of a temporal sequence. Within a temporal chunk, neural activity may show a sustained deviation from baseline (Barascud et al., 2016, Peña and Melloni, 2012) or a monotonic change in phase or power (O’Connell et al., 2012, Pallier et al., 2011, Brosch et al., 2011). At the end of a temporal sequence, an offset response is often observed (Ding et al., 2016, Nelson et al., 2017, Brennan et al., 2016). Furthermore, the sensory responses to individual events within a temporal chunk are significantly altered by learning (Gavornik and Bear, 2014, Sanders et al., 2002, Yin et al., 2008, Zhou et al., 2010, Farthouat et al., 2016, Buiatti et al., 2009), demonstrating that prior knowledge strongly influences sensory processing of sequences.

Whether sequential grouping requires attention is under debate (Snyder et al., 2006, Shinn-Cunningham, 2008, Shinn-Cunningham et al., 2017). On the one hand, it has been hypothesized that top-down attention is required for sequential grouping, especially for complex scenes consisting of multiple sequences. Evidence has been provided that attention can strongly affect neural and behavioral responses to sound sequences (Carlyon et al., 2001, Shamma et al., 2011, Lu et al., 2017, Fritz et al., 2007). Research on visual object recognition has also suggested that top-down attention is required for the binding of simultaneously presented features, e.g., color and shape information (Treisman and Gelade, 1980). On the other hand, a large number of neurophysiological studies have shown that the brain is highly sensitive to temporal regularities in sound when when the sound is not attended (Barascud et al., 2016, Näätänen et al., 2007, Sussman et al., 2007), suggesting that primitive analyses of temporal sequences may occur as a preattentative automatic process (Fodor, 1983).

Sequential grouping is not a single computational module, which further complicates the discussion about how attention influences sequential grouping. Sequential grouping can depend on multiple mechanisms, including bottom-up primitive grouping and top-down schema-based grouping (Bregman, 1990). Bottom-up grouping depends on the similarity between sensory features (Micheyl et al., 2005, McDermott et al., 2011, Woods and McDermott, 2015) while top-down schema-based grouping relies on prior knowledge (Billig et al., 2013, Hannemann et al., 2007, Jones and Freyman, 2012). Both grouping mechanisms play important roles in auditory perception. For example, in spoken word recognition, integrating acoustic features into phonemes and syllables can rely on acoustic continuity cues within a syllable (Shinn-Cunningham et al., 2017) while integrating syllables into words crucially relies on lexical knowledge, i.e., the knowledge about which syllable combinations constitute valid words (Cutler, 2012). Most previous studies focus on how attention modulates primitive sequential grouping while relatively little is known about how schema-based grouping is modulated by attention. The current study fills this gap by studying how the brain groups syllables into words based on lexical knowledge.

Behavioral evidence has suggested that cognitive processing of unattended spoken words is limited. Without paying attention, listeners cannot recall the spoken words they heard and cannot even notice a change in the language being played (Cherry, 1953). There is also evidence, however, for some low-level perceptual analysis for the unattended speech stream. For example, listeners can recall the gender of an unattended speaker (Cherry, 1953) and some listeners can notice their names in the unattended speech stream (Conway et al., 2001, Wood and Cowan, 1995). These results suggest that different speech processing stages could be differentially influenced by attention. Basic acoustic features can be recalled, very salient words such as one’s name can sometimes be recalled, while ordinary words cannot be recalled.

In this study, we used spoken word processing as a paradigm to test how attention may differentially modulate neural processing of basic sensory events, i.e., syllables, and temporal chunks constructed based on prior knowledge, i.e., multisyllabic words. Recent human neurophysiological results showed that cortical activity could concurrently follow hierarchical linguistic units of different sizes (Ding et al., 2016). In this study, we employed an isochronous syllable sequences as the speech stimulus, in which neighboring syllables combined into bisyllabic words (Fig. 1A). The stimulus was made in Chinese and all the syllables are monosyllabic morphemes. We first tested whether neural entrainment at the word rate could be observed, without any acoustic cue between word boundaries, and then tested whether attention differentially modulated neural entrainment to syllables (acoustic events) and neural entrainment to bisyllabic words (temporal chunks). The listener’s attentional focus was differently manipulated in three experiments. Experiment one and two presented competing sensory stimuli, e.g., a spoken passage or a silent movie, together with the isochronous syllable sequence, and the listeners had to attend to different stimuli in different experimental blocks. Experiment three, in contrast, directed the listener’s attentional focus to specific cued time intervals.

**Figure 1.**
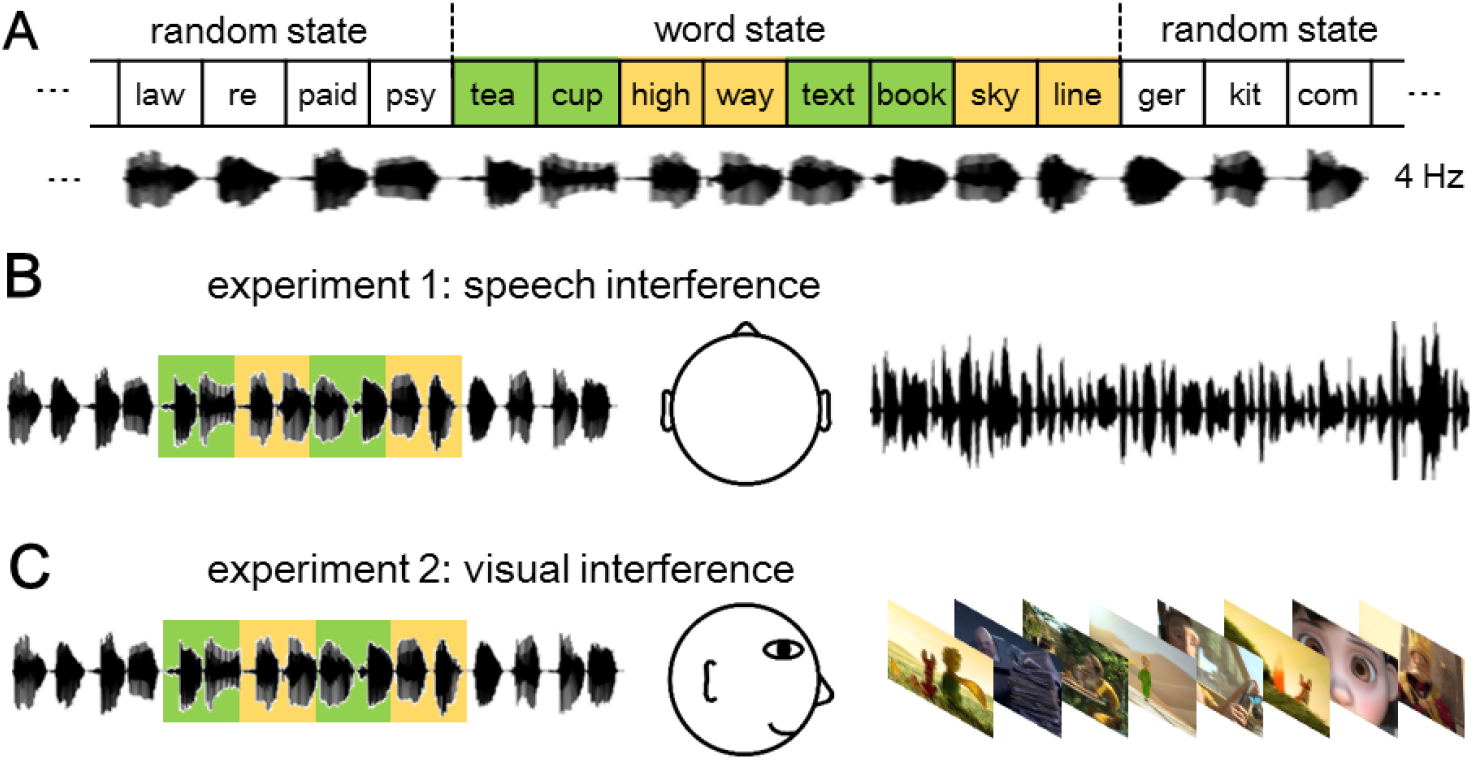
Experiment design. (A) Structure of the isochronous syllable sequence, which alternates between word states and random states. The syllables are presented at a constant rate of 4 Hz and therefore the bisyllabic words are presented at 2 Hz. English syllables are shown in the figure for illustrative purposes and Chinese syllables and words are used in the experiment. (B) In experiment one, the isochronous syllable sequence and a competing spoken passage are simultaneously presented to different ears. (C) In experiment two, the listeners either attend to the isochronous syllable sequence (presented to both ears) or watch a movie while passively listening to the syllable sequence.

## Results

In the first experiment, listeners were exposed to two concurrent speech streams, one to each ear (i.e., dichotically). One speech stream was an isochronous syllable sequence that alternates between word states and random states (Fig. 1). In the word states neighoring two words constructed a bisyllabic words and in the random state the order between syllables was randomized. The other speech stream was a spoken passage that was time compressed, i.e. fastened, by a factor of 2.5 to increase task difficulty.

During the time intervals when the bisyllabic words are played, the EEG power spectrum averaged over subjects and channels is shown in Fig. 2A. When the word sequence is attended, two peaks are observed in the power spectrum, one at the syllabic rate (P = 10^−4^, bootstrap) and the other at the word rate (P = 10^−4^, bootstrap). The topographic distribution of EEG power is centered near channel FCz. When attention is directed to the competing speech stream, a single response peak is observed at the syllabic rate (P = 10^−4^, bootstrap) while the neural response at the word-rate is no longer significantly stronger than the power in the neighboring frequency bins (P = 0.58, bootstrap). Comparing the conditions when the word lists are attended to or not, the difference in normalized word-rate response amplitude (i.e., the difference between the filled red and black bars in Fig. 2B) is significantly larger than the difference in normalized syllable-rate response amplitude (i.e., the difference between the hollow red and black bars in Fig. 2B, P = 0.01, bootstrap). The change in normalized word-rate response amplitude is more than 21.7 dB larger than the change in normalized syllable-rate response amplitude (27 dB vs. 5.3 dB). These results demonstrate that selective attention has a much stronger influence on the neural representation of linguistically defined temporal chunks, i.e., words, than the neural representation of acoustic events, i.e., syllables.

**Figure 2.**
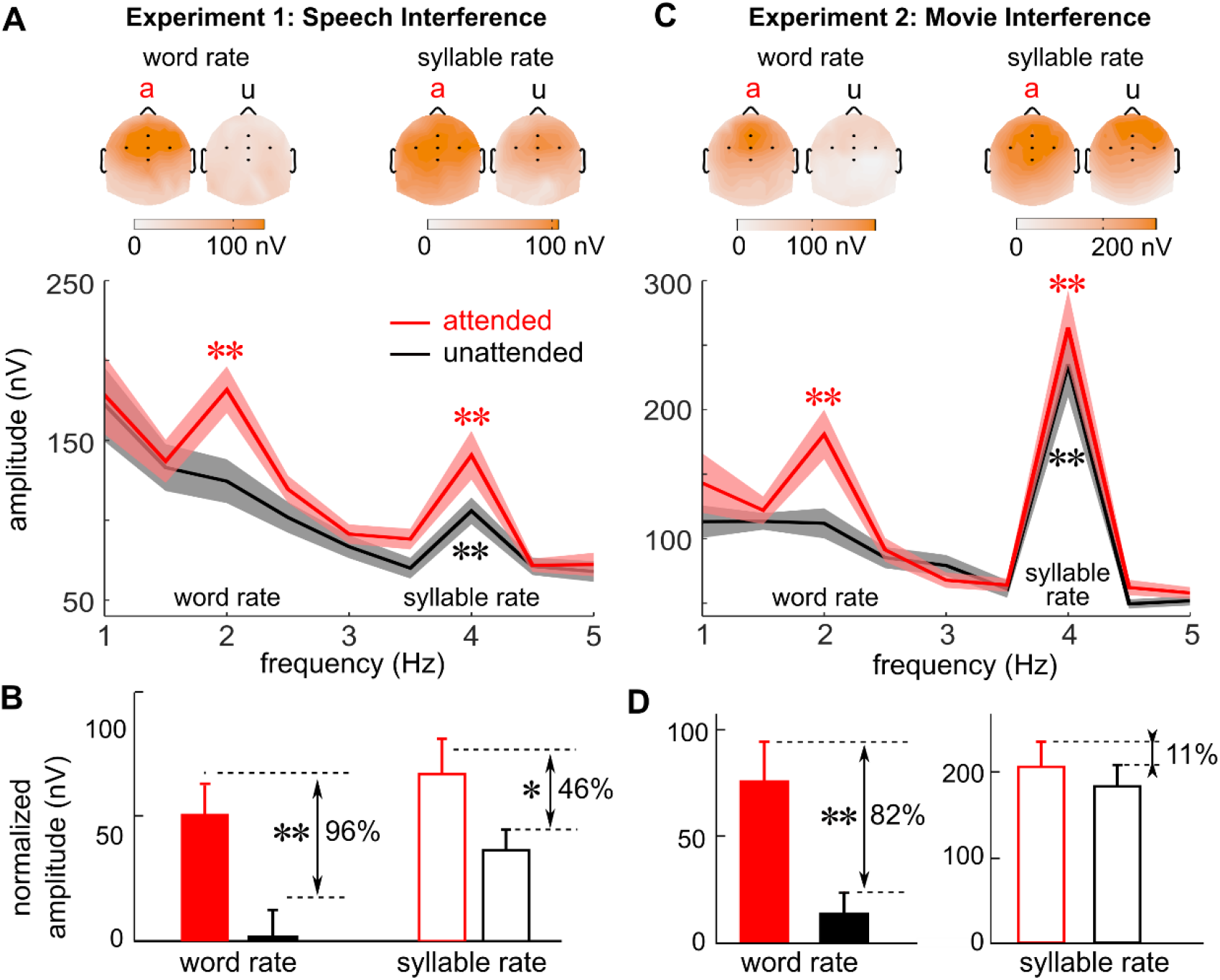
Attention differentially modulates neural entrainment to syllables and bisyllabic words. EEG response spectra averaged over subjects and channels are shown in panel A and C for experiment one and two respectively. Stars indicate frequency bins that show significantly stronger power than the power averaged over a 1-Hz wide neighboring frequency region (* P < 0.05, ** P < 0.005, bootstrap). Response peak at the syllabic rate is observed in both attended and unattended conditions. Response peak at the word rate however, is only observed for the attended condition. The topographic plots of the EEG response at the syllable and word rates are shown above the spectrum (a: attended; u: unattended), which generally shows a central-frontal distribution. In the topographic plots, the 5 black dots show the position of FCz (middle), Fz (upper), Cz (lower), FC3 (left), and FC4 (right). (B,D) Normalized power at the syllable and word rates. Power at each target frequency is normalized by subtracting the power averaged over a 1-Hz wide neighboring frequency region (excluding the target frequency), which reduces the influence of background broadband neural activity. Red bars represent the attended condition and black bars represent the unattended condition. The attention-related amplitude change relative to the response amplitude in the attended condition, i.e. (attended-unattended)/ attended, is shown in percentage near each response peak. Stars indicate whether the attention-related change in response amplitude is significantly larger than 0. Attention modulates both the syllable-rate response and the word-rate response but the effect is much stronger at the word rate.

Spoken passage comprehension involves almost all neural computations required for spoken word recognition. Therefore, it remains unclear whether the strong modulation of word-rate processing is due to the lack of top-down attention or a competition in other neural resources required for spoken word recognition. To address this issue, experiment 2 utilizes visual input to divert top-down attention. In this experiment, the isochronous syllable sequence is presented to both ears diotically and listeners either listen to speech or watch a silent movie with subtitles. The EEG power spectrum during the time intervals when the word states are presented is shown in Fig. 2C. The results largely mirror the results in experiment 1, except that the word-rate response is marginally significant when attention is directed to the visual input (P = 0.07, bootstrap). The attention related change in response amplitude is stronger at the word rate than at the syllable rate (i.e., the amplitude difference between the filled red and black bars in Fig. 2D is larger than the amplitude difference between the hollow red and black bars, P = 0.002, bootstrap). These results show that without any competing auditory input, the word-level neural representation still strongly relies on top-down attention.

The frequency-domain analysis in Fig. 2 reveals steady-state properties of the neural tracking of syllables and words. To further reveal how the neural response evolves over time, the waveform of the EEG signals averaged over channels is shown in Fig. 3. EEG responses show clear syllabic-rate oscillations when listening to random syllables. When bisyllabic words appear, EEG activity becomes dominated by word-rate oscillations, as is revealed by the intervals between response peaks (Fig. 3AB). The neural response power near the word and the syllable rates is further illustrated in Fig. 3CD. In the attended conditions, the word-rate neural response starts to increase about 500 ms after the word state onset when speech is presented in quiet (Fig. 3D) and this latency elongates to about 1 s when there is a competing speech stream (Fig. 3C). Furthermore, the syllabic-rate neural response shows a decrease in power about 1 s after the word state onset (Fig. 3CD).

**Figure 3.**
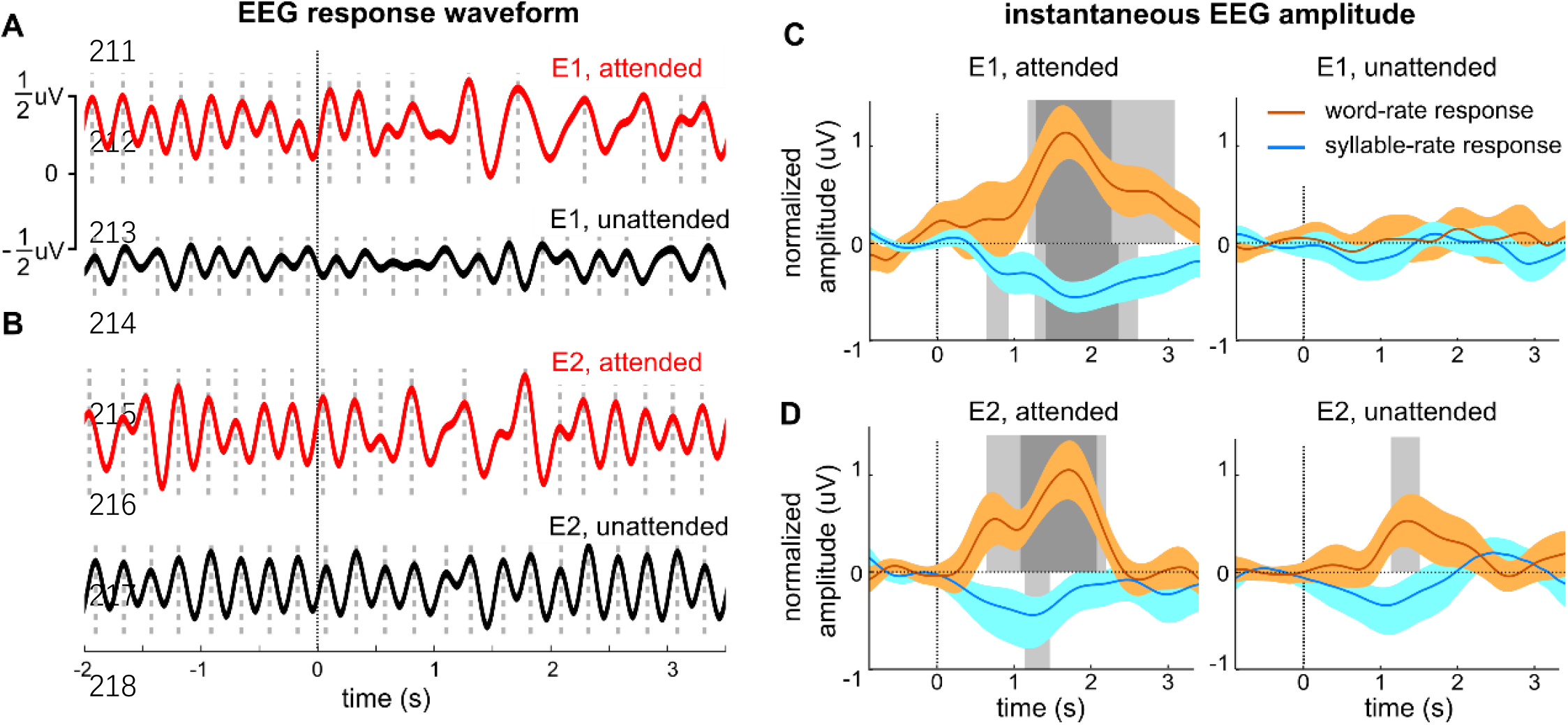
Temporal dynamics of the EEG response to words. (AB) The EEG waveforms for experiment one (E1) and experiment two (E2) are shown in panel A and B respectively (bandpass filtered between 1.5 and 4.5 Hz). The EEG waveform is grand averaged over subjects and channels. The word state starts from time 0. Each response peak, i.e., local maximum, is marked by a dotted line. Before the onset of the word state, regular neural oscillations are observed showing a peak every ∼250 ms, corresponding to a 4-Hz syllable-rate rhythm. About 500-1000 ms after the word onset, in attended conditions, a slow oscillation emerges showing a peak every 500 ms, corresponding to a 2-Hz word-rate rhythm. (CD) Instantaneous amplitude of the EEG response filtered around the syllable rate (1.75-2.25 Hz) or the word rate (3.75-4.25 Hz) for experiment one and two. The EEG instantaneous amplitude is baseline corrected by subtracting the mean amplitude in a 1-second duration pre-stimulus interval. The shaded areas above/below the horizontal dotted line at 0 μV indicate time intervals when the word-/syllable-rate response amplitude significantly differs from the pre-stimulus baseline (dark gray: P < 0.01, light gray: P<0.05; bootstrap, FDR corrected). The word-rate response shows a significant increase in power about 500-1000 ms after the first word appears, in all conditions except for the unattended condition in experiment one. For the attended conditions, a decrease in the syllable-rate response is also seen during the word state. The instantaneous amplitude is the magnitude of the Hilbert transform of the filtered EEG responses.

Experiments 1 and 2 show that neural tracking of words is severely attenuated when attention is directed to a competing sensory stimulus. We then ask if attention can modulate the word-rate neural response dynamically over time, in the absence of any competing stimulus. In a 3^rd^ experiment, the listeners hear a single speech stream and have to attend to some words while ignoring others. The onset of each word state is verbally cued and the listeners have to focus on either the first word or the last word in a word state (Fig. 4A).

**Figure 4.**
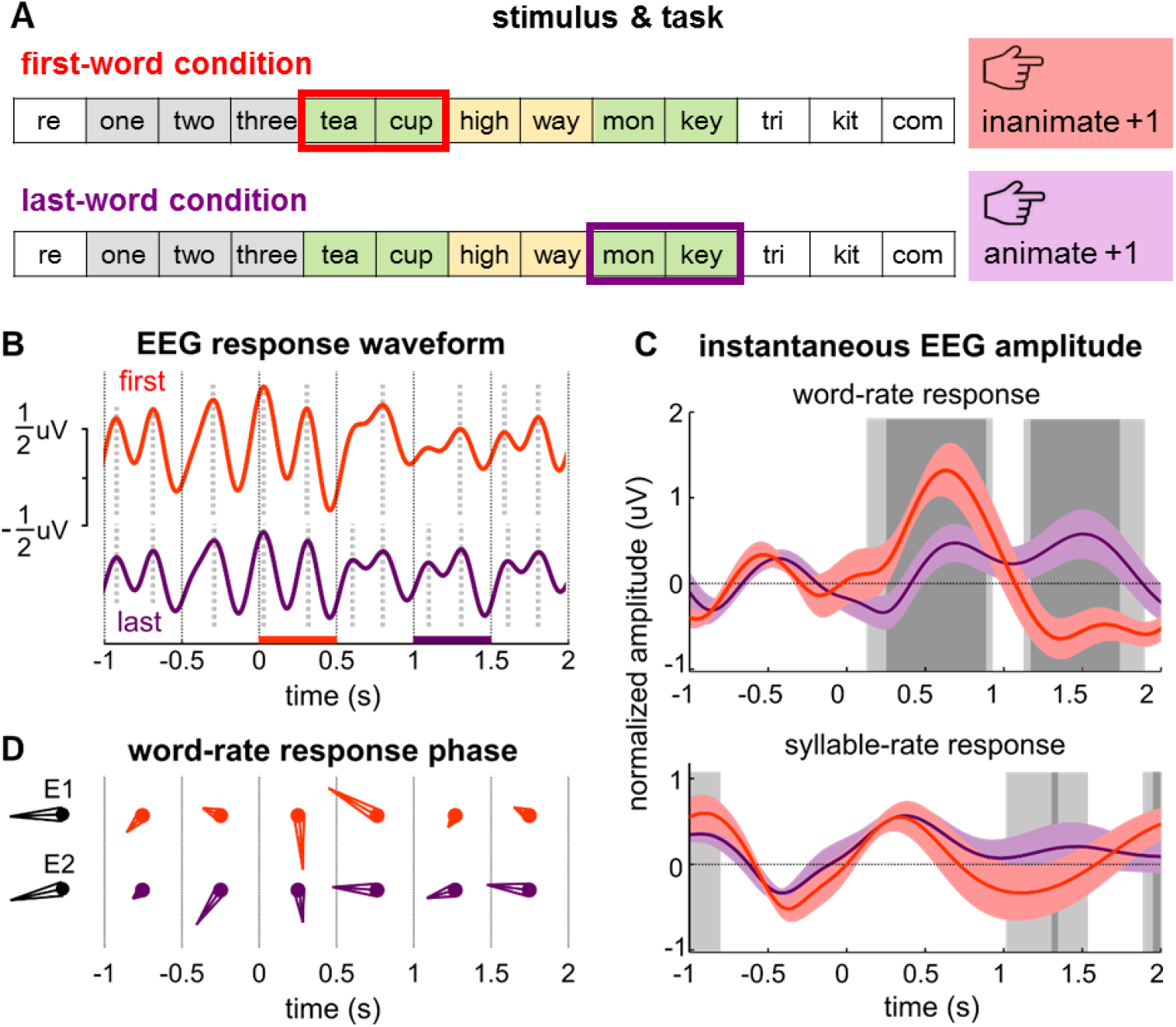
Temporal attention quickly modulates the neural tracking of words. (A) Illustration of the two tasks. The subjects have to judge the animacy of either the first word or the last word in a word state. The onset of the word state is cued by the preceding 3 syllables, which are one, two, and three. (B) The grand-averaged EEG response in the first-word condition and the last-word condition (using the convention in Fig. 3AB). The red and purple bars on the x-axis show the time intervals in which the first word and the last word in the word state are presented. (C) Instantaneous amplitude of the EEG response filtered around the syllable rate (1.75-2.25 Hz) and the word rate (3.75-4.25 Hz). The shaded areas indicate time intervals when the response amplitude significantly differs from the pre-stimulus baseline (dark gray: P < 0.01, light gray: P<0.05; bootstrap, FDR corrected). The word-rate response is strongly modulated by temporal attention and shows stronger activation near the attended word (i.e., stronger activation in an earlier window for the first-word condition compared with the last-word condition). The syllable-rate response is less strongly modulated by temporal attention. The instantaneous amplitude is extracted using the same method used in Fig. 3CD. (D) Phase and amplitude of the word-rate response in each 500-ms time bin. The red and purple arrows indicate complex-valued Fourier coefficient at the word rate in each time bin, for the first- and last-word conditions respectively. The response shows a phase change between the first bin and the second bin in the word state. The black arrows show the mean response averaged over the word state response in experiment one (E1) and experiment two (E2).

In experiment 3, the listeners have to judge the animacy of the first word in each word state in one block (called the first-word condition) and judge the animacy of the last word in each word state in another block (called the last-word condition). Timing is critical for these tasks since the listeners have to judge the animacy of the right words and not confuse them with the neighboring words. The two tasks force the listeners to attend to words at different positions of a sequence and therefore dissociate their attentional focus in time.

The results of experiment 3 are shown in Fig. 4. The time course of the word-rate neural response is significantly modulated by temporal attention. The neural response shows a stronger word-rate response near the beginning/end of a word state in the first/last word condition (Fig. 4B). In other words, the word-rate response is significantly stronger during the time intervals being attended to. Although the onset of the word state is cued, the phase of the word-rate response still takes about 500 ms to stabilize after the word state onset (Fig. 4C). In other words, temporal prediction cannot greatly fasten the stabilization of the neural response phase.

## Discussion

The current study investigates how attention differentially modulates the neural entrainment to acoustic events, i.e., syllables, and temporal chunks, i.e., words. Here, the grouping of syllables into words is purely based on top-down lexical knowledge, i.e., the mental dictionary, rather than bottom-up acoustic cues. It is shown that top-down attention more strongly modulates the word-rate neural response compared with the syllable-rate neural response (up to 20 dB differences in attention-related changes in response power), which strongly suggests that attention is crucial for knowledge-based sequential grouping.

### Neural processing of unattended auditory streams

The brain can detect statistical regularities in sounds even without top-down attentional modulation (Näätänen et al., 2007). For example, neural activity can entrain to intensity fluctuations in sound even when the listeners do not pay attention (Linden et al., 1987). Similarly, in the current study, the syllabic rhythm is reflected in the EEG response whether the listeners pay attention or not. Previous studies have shown that when a random tone cloud turns into a fixed multi-tone sequence repeating in time, the brain can quickly detect such a transition even when attention is directed to other sensory stimuli (Barascud et al., 2016). Furthermore, the brain can also detect violations in multi-tone sequences that repeat in time (Sussman et al., 2007). Therefore, although attention can strongly modulate primitive auditory grouping, i.e., bottom-up feature-based grouping of acoustic events into auditory streams (Carlyon et al., 2001, Shamma et al., 2011, Shinn-Cunningham et al., 2017), it is clear that the brain can detect basic statistical regularities in sounds preattentatively.

Statistical regularities in sound can be extracted by bottom-up analysis of auditory features. In the current study, however, the grouping of syllables into words can only rely on top-down knowledge about which syllables can possibly construct a valid multisyllabic word. The word boundaries can only be determined by comparing the auditory input with word templates stored in the long-term memory. The current results show that neural entrainment to bisyllablic words is much more strongly influenced by top-down attention, compared with the neural entrainment to syllables. Therefore, although bottom-up grouping of basic auditory features into a sound stream may occur preattentatively, top-down schema-based grouping of syllables into words critically relies on attention.

### Attention modulation of neural processing of speech

This study uses Chinese as the testing language. In Chinese, generally speaking, each syllable corresponds to a morpheme but there is no one-to-one mapping between syllables and morphemes due to the existence of homophones. For example, the syllable 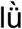 could correspond to an adjective (e.g., green 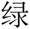), a noun (e.g., law 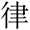), or a verb (e.g., filter 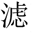). Since the mapping between syllables and morphemes is highly ambiguous, a random syllable sequence cannot be reliably mapped into a sequence of morphemes and is generally heard as a meaningless syllable sequence. The bisyllabic words used in this study, however, are common unambiguous words that can be precisely decoded when listening to the syllable sequences. Therefore, the study probes the process of grouping syllables (ambiguous morphemes) into multisyllabic (multimorphemic) words.

Speech comprehension involves multiple processing stages, e.g., encoding acoustic speech features (Shamma, 2001), decoding phonemic information based on acoustic features (Mesgarani et al., 2014, Di Liberto et al., 2015), grouping syllables into words (Cutler, 2012), and grouping words into higher level linguistic structures such as phrases and sentences (Friederici, 2002). Previous studies have shown that attention can modulate neural entrainment to the intensity fluctuations in the speech, i.e., the speech envelope that corresponds to the syllabic rhythm (Kerlin et al., 2010, Mesgarani and Chang, 2012, O’Sullivan et al., 2014, Park et al., 2016). The envelope-following response is stronger for the attended speech but remains observable for the unattended speech (Ding and Simon, 2012, Steinschneider et al., 2013), especially when there is no competing auditory input (Kong et al., 2014). In terms of the spatial distribution of neural activity, neural entrainment to the unattended speech is stronger near sensory areas around the superior temporal gyrus and attenuates in higher-order cortical areas (Golumbic et al., 2013). The current study extends previous studies by showing that neural entrainment to linguistic units, such as words, is more strongly modulated by attention than neural entrainment to the speech envelope. When attention is directed to a competing speech stream, word-rate neural entrainment is no longer observed. These results show that attention strongly modulates the lexical segmentation process, which creates a bottleneck for the neural processing of unattended speech streams.

Previous studies on attention modulation of lexical processing mostly focus on semantic processing of words that have clear physical boundaries. It is found that the N400 ERP response disappears for unattended auditory or visual words (Nobre and Mccarthy, 1995, Bentin et al., 1995). On the other hand, visual experiments have shown that semantic processing can occur for words presented at the attended location even when these words are not consciously perceived (Luck et al., 1996, Naccache et al., 2002). Therefore, semantic processing of isolated words could be a subconscious process but requires attention. The current study extends these previous studies by showing the phonological construction of words, i.e., the grouping of syllables into words, also requires attention. Here, the grouping of syllables into words can only be achieved by comparing the input speech stream with phonological templates of words that are stored in long-term memory. Therefore, the current results strongly suggest that phonological grouping process crucially relies on attention.

### Low-frequency neural oscillations and temporal information processing

The current data and previous studies (Ding et al., 2016, Buiatti et al., 2009, Steinhauer et al., 1999, Farthouat et al., 2016, Meyer et al., 2016, Peelle et al., 2013) show that, during speech listening, cortical activity is concurrently entrained to hierarchical linguistic units, including syllables, words, phrases, and sentences. Neural entrainment to hierarchical linguistic units provides a plausible mechanism to map hierarchical linguistic units into coupled dynamic neural processes that allow interactions between different linguistic levels (Martin and Doumas, 2017, Goswami and Leong, 2013, Giraud and Poeppel, 2012, Wassenhove et al., 2003). Neural entrainment to words stabilizes ∼0.5-1 s after the word state onset. Similarly, previous studies have shown that neural entrainment to phrases and sentences also stabilizes within ∼1 ms (Zhang and Ding, 2017). When the onset time of a word state is precisely cued, the neural response phase still takes about ∼0.5 s to stabilize (Fig. 4C), suggesting that neural entrainment to words is not purely a predictive process and requires feedfowrd syllabic input.

The current data and previous results (Ding et al., 2016) suggest that low-frequency neural entrainment is closely related to the binding of syllables into temporal chunks such as words and phrases. Previous studies have also suggested slow changes in neural activity may indicate information integration over time during word by word reading (Pallier et al., 2011) and during decision making (O’Connell et al., 2012). Therefore, low-frequency neural entrainment provides a plausible neural signature for the mental construction of temporal chunks.

Low-frequency neural entrainment to sensory stimuli is a widely observed phenomenon. Neurophysiological evidence has been provided that the phase of low-frequency neural oscillations can modulate neuronal firing (Lakatos et al., 2005, Canolty et al., 2006) and can serve as a mechanism for temporal attention and temporal prediction(Arnal and Giraud, 2012, Schroeder and Lakatos, 2009). Furthermore, slow neural oscillations may also provide a neural context for the integration of faster neural activity falling into the same cycle of a slow neural oscillation (Buzsáki, 2010, Lisman and Jensen, 2013). Therefore low-frequency neural entrainment to temporal chunks may naturally provide a mechanism to put neural representations of sensory events into a context and allow information integration across sensory events.

## Methods

### Subjects

Fourteen subjects participated in each experiment (18-28 years old; mean age: 22; 50% female). All subjects were graduate or undergraduate students at Zhejiang University, with no self-reported hearing loss or neurological disorders. The experimental procedures were approved by the Institutional Review Board of Zhejiang University Interdisciplinary Center for Social Sciences. The subjects provided written consent and were paid for the experiment.

### Word Materials

The study employed 160 animate bisyllabic words and 160 inanimate bisyllabic words. Animate words included animals (N = 40, e.g., monkey, dolphin), plants (N = 40, e.g. lemon, carrot), humans (N = 48, e.g., doctor, doorman), and names of well known people in history (N = 32, e.g., Bai Li, a famous poet in Tang dynasty). Inanimate words include objects (N = 80, e.g., teacup, pencil) and places (N = 80, e.g., Beijing, Zhejiang).

### Stimuli

The stimulus consisted of an isochronous syllable sequence. All syllables were independently synthesized using the Neospeech synthesizer (http://www.neospeech.com/, the male voice, Liang). All syllables were adjusted to the same intensity and the same duration, i.e., 250 ms (see Ding et al., 2016 for details). The syllable sequence alternated between a word state and a random state (Fig. 5A). The number of syllables in each state and the number of word states in each stimulus, i.e., *M*, were shown in Fig 5B. Each sequence started and ended with a random state to reduce the probability that words might pop out at the beginning and end of each stimulus, even when the syllable sequence was not attended.

**Figure 5.**
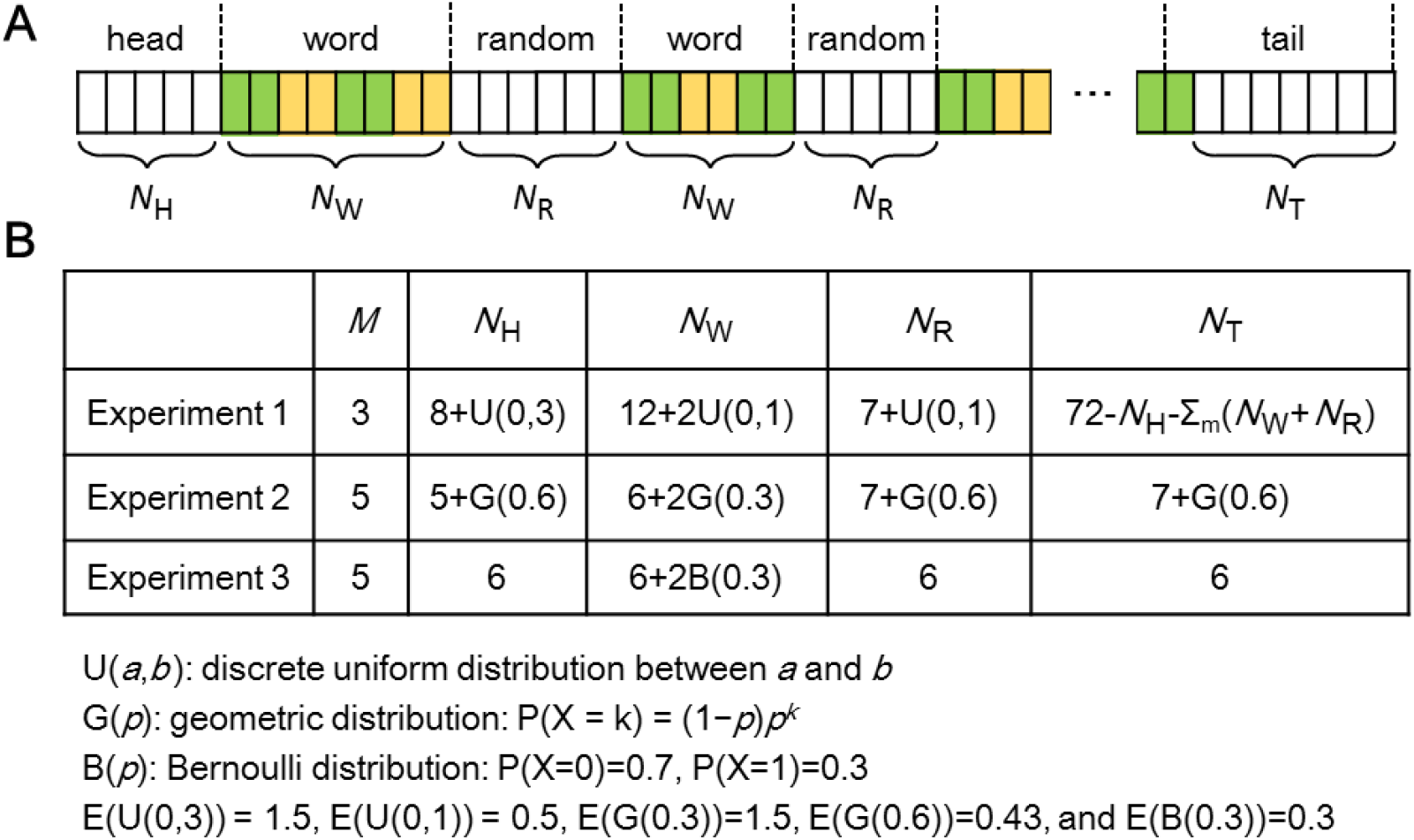
Structure of the isochronous syllable sequence in each experiment. (A) The sequence alternates between random states and word states M times in each trial. At the beginning and end of each trial, NH and NT random syllables are presented. (B) Statistical distribution of the number of syllables in each state.

In experiment one, an isochronous syllable sequence and a competing spoken passage were dichotically presented, and the ear each stimulus was presented to was counterbalanced across subjects. The competing spoken passages (chosen from the *Syllabus for Mandarin Proficiency Tests*) were time compressed by a factor of 2.5 and gaps longer than 30 ms were shortened to 30 ms. Long acoustic pauses were removed in case the listeners might shift their attentional focus during the pauses. In each trial, 19 seconds of spoken passages were presented and the duration of each syllable sequence was set to 18 seconds, i.e., 72 syllables. The competing spoken passage started 1 second before the syllable sequence so that the syllable sequence was less likely to be noticed when the listeners focused on the spoken passage. The number of syllables in the word and random states was randomized using a uniform distribution so that the alternation between states was not completely regular while the total duration could be easily controlled.

In experiment two, an isochronous syllable sequence was identically, i.e., diotically, presented to both ears. The number of syllables in the word and random states was subject to a geometric distribution so that the subjects could not predict when state transitions would occur.

In experiment three, each random state always consisted of 6 syllables and the last 3 syllables were always “yi, er, san” which means “one two three” in mandarin Chinese. These 3 syllables served as cues for the onset time of a word state.

In all experiments, no word appeared twice in a trial and there was no immediate repetition of any syllable. In experiment one and two, words in the same word state belonged to the same category, i.e., animate or inanimate. In experiment three, however, the words in each word state were randomly chosen from all possible words. The subjects were never told how many word states might appear in a trial.

### Procedures

The study consisted of three experiments. Each experiment contained two blocks, differing in the subject’s attentional focus.

**Experiment one:** In the first block, listeners had to focus on the time-compressed spoken passage and answer comprehension questions after each trial. The comprehension questions were presented 1 s after the spoken passage and the listeners had to give a verbal answer (correct rate: 84 ± 2%, mean ± standard error throughout the paper). After the experimenter recorded the answer they pressed a key to continue the experiment. The next trial was played after an interval randomized between 1 and 2 seconds (uniform distribution) after the key press. In the second block, subjects had to focus on the syllable sequences and judge if an additional word presented 1 s after the sequence offset appeared in the sequence by a key press (correct rate: 77 ± 2%). The next trial started after an interval randomized between 1 and 2 seconds (uniform distribution) after the key press. The same set of 50 trials (50 distinct spoken passages paired with 50 distinct syllable sequences) were presented in each block with a random order. The subjects had their eyes closed when listening to the stimuli and had a break every 25 trials. The listeners always attended to the spoken passages in the first block to reduce the possibility that they may spontaneously shift their attentional focus to the isochronous syllable sequence after knowing that there were words embedded in the sequence.

**Experiment two**: A word listening block and a movie watching block were presented, the order of which was counterbalanced across subjects. In the word listening block, after each trial, the subjects had to judge if they heard more animate words or more inanimate words by pressing different keys (correct rate: 81±3%). The subjects were told that all words within the same word state belonged to the same category, i.e., animate or inanimate. Sixty trials were presented and the subjects had a break after every 15 trials. Before the word listening condition, the subjects went through a practice section, in which they listened to two example sequences and did the same task. They received feedback during the practice session but not during the main experiment. The neural responses showed the same pattern whichever block was presented first and therefore the responses were averaged over all subjects regardless of the presentation order.

In the movie watching block, the subjects watched a silent movie (the Little Prince) with Chinese subtitles. The syllable sequences were presented about 3 minutes after the movie started to make sure that the subjects had already engaged in the movie watching task. Sixty syllable sequences were presented in a randomized order, with the inter-stimulus-interval randomized between 1 and 2 seconds. The movie was stopped after all the 60 sequences were presented. The subjects had their eyes open in both blocks although no visual stimulus was presented in the word listening block.

**Experiment three:** The experiment was divided into a first-word condition block and a last-word condition block, the order of which were counterbalanced across subjects. The subjects had to judge whether they heard more animate words or inanimate words by pressing different keys. In the first-word/last-word condition, they should only count the first-word or the last word in each word state. Five word states appeared in each trial and therefore if, e.g., 3 word states started with animate words the subjects should judge that the trial had more animate words in the first-word condition. They were not told how many word states might appear in each sequence. The subjects had a break every 15 trials. Before each condition, the subjects went through a practice session, in which they listened to two example sequences and made judgments. They received feedback during the practice session but not during the main experiment. In the main experiment, the subjects gave correct answers in 80 ± 4% and 63 ± 3% trials in the first-word and last-word conditions respectively. The correct rate was significantly higher in the first-word condition (P < 0.0001, bootstrap), in which the timing of the target word was more predictable. The correct rate, however, remained above the 50% chance level in the last-word condition (P < 0.0001, bootstrap).

### EEG recording and analysis

EEG responses were recorded using a 64-channel Biosemi ActiveTwo system. Additionally, four electrodes were used to record horizontal and vertical EOG and two reference electrodes were placed at the left and right mastoids. The EEG recordings were low-pass filtered below 400 Hz and sampled at 2048 Hz. The EEG recordings were referenced to the average mastoid recording offline and the horizontal and vertical EOG signals were regressed out. Since the study focused on word-rate and syllable-rate neural responses (2 Hz and 4 Hz respectively), the EEG recordings were high-pass filtered above 0.7 Hz. The EEG recordings were epoched based on the onset of each word state (9 s epochs starting 2 s before the word state onset) and averaged over all epochs.

In the frequency domain analysis, a Discrete Fourier Transform was applied to each EEG channel and each subject. The analysis window was 2 s in duration, corresponding to a frequency resolution of 0.5 Hz. In experiment two, a single analysis window was used, which started from the word state onset. In experiment one, since the word state is longer, two successive analysis windows were applied, with the first one starting from the word state onset and the second starting from the offset of the first analysis window. The EEG spectrum is averaged over EEG channels and subjects (and also analysis windows in experiment one) by calculating the root-mean-square value.

In the time domain analysis, to visualize the response waveform (Fig. 3A), the EEG responses were filtered between 1.5 and 4.5 Hz using a linear phase finite impulse response (FIR) filter (impulse response duration: 1 s). The linear delay caused by the FIR filter is compensated by shifting the filtered signal back in time. When separately analyzing the instantaneous amplitude of the word-rate or syllable-rate response (Fig. 3CD and 4C), the EEG responses were bandpass filtered using a 1-s duration FIR filter with the lower and higher cutoff frequencies set to 0.25 Hz below and above the word or syllable rate. The instantaneous amplitude of the word-rate and syllable-rate EEG responses were extracted using the Hilbert transform.

### Statistical test

This study used bias-corrected and accelerated bootstrap for all significance tests (Efron and Tibshirani, 1993). In the bootstrap procedure, all the subjects were resampled with replacement 10^4^ times. For the significance test for peaks in the response spectrum (Fig. 2A), the response amplitude at the peak frequency is compared with the mean amplitude of the neighboring 2 frequency bins (corresponding to a 1-Hz width). For the significance test for time intervals showing response amplitude differences (Fig. 3CD and 4C), the EEG waveform was averaged over all sampled subjects and the instantaneous amplitude was then extracted using the Hilbert transform.

## Acknowledgement

We thank Huan Luo, Lucia Melloni, David Poeppel, Jonathan Simon, and Elana Zion-Golumbic for helpful comments on earlier versions of this manuscript.

### Funding

Work supported by National Natural Science Foundation of China 31500873 (ND), Zhejiang Provincial Natural Science Foundation of China LR16C090002 (ND), Fundamental Research Funds for the Central Universities (ND), and research funding from the State Key Laboratory of Industrial Control Technology, Zhejiang University (ND). The funding sources were not involved in study design, data collection and interpretation, or the decision to submit the work for publication.

## References

Arnal, L. H. & Giraud, A.-L. 2012. Cortical oscillations and sensory predictions. Trends in Cognitive Sciences, 16, 390–398.

Barascud, N., Pearce, M. T., Griffiths, T. D., Friston, K. J. & Chait, M. 2016. Brain responses in humans reveal ideal observer-like sensitivity to complex acoustic patterns. Proceedings of the National Academy of Sciences, 113, E616–E625.

Bentin, S., Kutas, M. & Hillyard, S. A. 1995. Semantic processing and memory for attended and unattended words in dichotic listening: behavioral and electrophysiological evidence. Journal of Experimental Psychology: Human Perception and Performance, 21, 54.

Billig, A. J., Davis, M. H., Deeks, J. M., Monstrey, J. & Carlyon, R. P. 2013. Lexical influences on auditory streaming. Current Biology, 23, 1585–1589.

Bregman, A. S. 1990. Auditory scene analysis: the perceptual organization of sound, Cambridge, The MIT Press.

Brennan, J. R., Stabler, E. P., Van Wagenen, S. E., Luh, W. M. & Hale, J. T. 2016. Abstract linguistic structure correlates with temporal activity during naturalistic comprehension. Brain and language, 157, 81–94.

Brosch, M., Selezneva, E. & Scheich, H. 2011. Formation of associations in auditory cortex by slow changes of tonic firing. Hearing research, 271, 66–73.

Buiatti, M., Pe A M. & Dehaene-Lambertz, G. 2009. Investigating the neural correlates of continuous speech computation with frequency-tagged neuroelectric responses. Neuroimage, 44, 509–51.

Buzsaki, G. 2010. Neural syntax: cell assemblies, synapsembles, and readers. Neuron, 68, 362–385.

Canolty, R. T., Edwards, E., Dalal, S. S., Soltani, M., Nagarajan, S. S., Kirsch, H. E., Berger, M. S., Barbaro, N. M. & Knight, R. T. 2006. High Gamma Power Is Phase-Locked to Theta Oscillations in Human Neocortex. Science, 313, 1626 – 1628.

Carlyon, R. P., Cusack, R., Foxton, J. M. & Robertson, I. H. 2001. Effects of attention and unilateral neglect on auditory stream segregation. Journal of Experimental Psychology: Human Perception and Performance, 27, 115–127.

Cherry, E. C. 1953. Some experiments on the recognition of speech, with one and with two ears. Journal of the Acoustical Society of America, 25, 975–979.

Conway, A. R. A., Cowan, N. & Bunting, M. F. 2001. The cocktail party phenomenon revisited: The importance of working memory capacity. Psychonomic Bulletin & Review, 8, 331–335.

Cutler, A. 2012. Native listening: Language experience and the recognition of spoken words, Mit Press.

Di Liberto, G. M., O'Sullivan, J. A. & Lalor, E. C. 2015. Low-frequency cortical entrainment to speech reflects phoneme-level processing. Current Biology, 25, 2457–2465.

Ding, N., Melloni, L., Zhang, H., Tian, X. & Poeppel, D. 2016. Cortical tracking of hierarchical linguistic structures in connected speech. Nature Neuroscience, 19, 158–164.

Ding, N. & Simon, J. Z. 2012. Emergence of neural encoding of auditory objects while listening to competing speakers. Proceedings of the National Academy of Sciences of the United States of America, 109, 11854–11859.

Efron, B. & Tibshirani, R. 1993. An introduction to the bootstrap, CRC press.

Farthouat, J., Franco, A., Mary, A., Delpouve, J., Wens, V., De Beeck, M. O., DE TI Ge, X. & Peigneux, P. 2016. Auditory Magnetoencephalographic Frequency-Tagged Responses Mirror the Ongoing Segmentation Processes Underlying Statistical Learning. Brain Topography, 1–13.

Fodor, J. A. 1983. The modularity of mind: An essay on faculty psychology, MIT press.

Friederici, A. D. 2002. Towards a neural basis of auditory sentence processing. Trends in cognitive sciences, 6, 78–84.

Fritz, J. B., Elhilali, M., David, S. V. & Shamma, S. A. 2007. Does attention play a role in dynamic receptive field adaptation to changing acoustic salience in A1? Hearing Research, 229, 186-203.

Gavornik, J. P. & Bear, M. F. 2014. Learned spatiotemporal sequence recognition and prediction in primary visual cortex. Nature neuroscience, 17, 732-737.

Giraud, A.-L. & Poeppel, D. 2012. Cortical oscillations and speech processing: emerging computational principles and operations. Nature neuroscience, 15, 511–517.

Golumbic, E. M. Z., Ding, N., Bickel, S., Lakatos, P., Schevon, C. A., Mckhann, G. M., Goodman, R. R., Emerson, R., Mehta, A. D., Simon, J. Z., Poeppel, D. & Schroeder, C. E. 2013. Mechanisms Underlying Selective Neuronal Tracking of Attended Speech at a “Cocktail Party". Neuron, 77, 980–991.

Goswami, U. & Leong, V. 2013. Speech rhythm and temporal structure: Converging perspectives? Laboratory Phonology, 4, 67–92.

Hannemann, R., Obleser, J. & Eulitz, C. 2007. Top-down knowledge supports the retrieval of lexical information from degraded speech. Brain research, 1153, 134–143.

Jones, J. A. & Freyman, R. L. 2012. Effect of priming on energetic and informational masking in a same-different task. Ear and hearing, 33, 124–133.

Kerlin, J. R., Shahin, A. J. & Miller, L. M. 2010. Attentional Gain Control of Ongoing Cortical Speech Representations in a “Cocktail Party”. Journal of Neuroscience, 30, 620–628.

Kong, Y.-Y., Mullangi, A. & Ding, N. 2014. Differential Modulation of Auditory Responses to Attended and Unattended Speech in Different Listening Conditions. Hearing Research, 316, 73–81.

Lakatos, P., Shah, A. S., Knuth, K. H., Ulbert, I., Karmos, G. & Schroeder, C. E. 2005. An oscillatory hierarchy controlling neuronal excitability and stimulus processing in the auditory cortex. Journal of neurophysiology, 94, 1904–1911.

Lashley, K. S. 1951. the problem of serial order in behavior. In: Jeffress, L. A. (ed.) Cerebral Mechanisms in Behavior, The Hixon Symposium. New York: Wiley.

Linden, R. D., Picton, T. W., Hamel, G. & Campbell, K. B. 1987. Human auditory steady-state evoked potentials during selective attention. Electroencephalography and Clinical Neurophysiology, 66, 145–159.

Lisman, J. E. & Jensen, O. 2013. The theta-gamma neural code. Neuron, 77, 1002–1016.

Lu, K., Xu, Y., Yin, P., Oxenham, A. J., Fritz, J. B. & Shamma, S. A. 2017. Temporal coherence structure rapidly shapes neuronal interactions. Nature Communications, 8, 13900.

Luck, S. J., Vogel, E. K. & Shapiro, K. L. 1996. Word meanings can be accessed but not reported during the attentional blink. Nature, 383, 616.

Martin, A. E. & Doumas, L. A. 2017. A mechanism for the cortical computation of hierarchical linguistic structure. PLoS Biology, 15, e200066.

Mcdermott, J. H., Wrobleski, D. & Oxenham, A. J. 2011. Recovering sound sources from embedded repetition. proceedings of the National Academy of Sciences, 108, 1188–1193.

Mesgarani, N. & Chang, E. F. 2012. Selective cortical representation of attended speaker in multi-talker speech perception. Nature, 485, 233–236.

Mesgarani, N., Cheung, C., Johnson, K. & Chang, E. F. 2014. Phonetic feature encoding in human superior temporal gyrus. Science, 343, 1006–1010.

Meyer, L., Henry, M. J., Gaston, P., Schmuck, N. & Friederici, A. D. 2016. Linguistic bias modulates interpretation of speech via neural delta-band oscillations. Cerebral Cortex.

Micheyl, C., Tian, B., Carlyon, R. P. & Rauschecker, J. P. 2005. Perceptual organization of tone sequences in the auditory cortex of awake macaques. Neuron, 48, 139–148.

Naatanen, R., Paavilainen, P., Rinne, T. & Alho, K. 2007. The mismatch negativity (MMN) in basic research of central auditory processing: a review. Clinical Neurophysiology, 118, 2544-2590.

Naccache, L., Blandin, E. & Dehaene, S. 2002. Unconscious masked priming depends on temporal attention. Psychological science, 13, 416-424.

Nelson, M. J., El Karoui, I., Giber, K., Yang, X., Cohen, L., Koopman, H., Dehaene, S. &. PROCEEDINGS OF THE NATIONAL ACADEMY OF SCIENCES 2017. Neurophysiological dynamics of phrase-structure building during sentence processing. Proceedings of the National Academy of Sciences, 114, E3669–E3678.

Nobre, A. C. & Mccarthy, G. 1995. Language-related field potentials in the anterior-medial temporal lobe: II. Effects of word type and semantic priming. Journal of neuroscience, 15, 1090–1098.

O'Connell, R. G., Dockree, P. M. & Kelly, S. P. 2012. A supramodal accumulation-to-bound signal that determines perceptual decisions in humans. Nature neuroscience, 15, 1729–1735.

O'Sullivan, J. A., Power, A. J., Mesgarani, N., Rajaram, S., Foxe, J. J., Shinn-Cunningham, B. G., Slaney, M., Shamma, S. A. & Lalor, E. C. 2014. Attentional Selection in a Cocktail Party Environment Can Be Decoded from Single-Trial EEG. Cerebral Cortex.

Pallier, C., Devauchelle, A.-D. & Dehaene, S. 2011. Cortical representation of the constituent structure of sentences. Proceedings of the National Academy of Sciences, 108, 2522–2527.

Park, H., Kayser, C., Thut, G. & Gross, J. 2016. Lip movements entrain the observers' low-frequency brain oscillations to facilitate speech intelligibility. eLife, 5, e14521.

Pena, M. & Melloni, L. 2012. Brain oscillations during spoken sentence processing. Journal of cognitive neuroscience, 24, 1149-1164.

Peelle, J. E., Gross, J. & Davis, M. H. 2013. Phase-Locked Responses to Speech in Human Auditory Cortex are Enhanced During Comprehension Cerebral Cortex, 23, 1378–1387.

Sanders, L. D., Newport, E. L. & Neville, H. J. 2002. Segmenting nonsense: an event-related potential index of perceived onsets in continuous speech. Nature neuroscience, 5, 700–703.

Schroeder, C. E. & Lakatos, P. 2009. Low-frequency neuronal oscillations as instruments of sensory selection. Trends in Neurosciences, 32, 9–18.

Shamma, S. 2001. On the role of space and time in auditory processing. Trends in cognitive sciences, 5, 340–348.

Shamma, S. A., Elhilali, M. & Micheyl, C. 2011. Temporal coherence and attention in auditory scene analysis. Trends in Neurosciences, 34, 114–123.

Shinn-Cunningham, B., Best, V. & Lee, A. K. 2017. Auditory Object Formation and Selection. *In:* Middlebrooks, J. C., Simon, J. Z., Popper, A. N. & Fay, R. R. (eds.) The Auditory System at the Cocktail Party. Springer International Publishing.

Shinn-Cunningham, B. G. 2008. Object-based auditory and visual attention. Trends in Cognitive Sciences, 12, 182–186.

Snyder, J. S., Alain, C. & Picton, T. W. 2006. Effects of attention on neuroelectric correlates of auditory stream segregation. Journal of cognitive neuroscience, 18, 1–13.

Steinhauer, K., Alter, K. & Friederici, A. D. 1999. Brain potentials indicate immediate use of prosodic cues in natural speech processing. Nature neuroscience, 2, 191–196.

Steinschneider, M., Nourski, K. V. & Fishman, Y. I. 2013. Representation of speech in human auditory cortex: Is it special? Hearing research, 57–73.

Sussman, E. S., HORV Th, J., Winkler, I. & Orr, M. 2007. The role of attention in the formation of auditory streams. Attention, Perception, & Psychophysics, 69, 136–152.

Treisman, A. M. & Gelade, G. 1980. A feature-integration theory of attention. Cognitive psychology, 12, 97–136.

Wassenhove, V. V., Grant, K. W. & Poeppel, D. 2003. Visual speech speeds up the neural processing of auditory speech. proceedings of the National Academy of Sciences of the United States of America, 102, 1181–1186.

Wood, N. & Cowan, N. 1995. The cocktail party phenomenon revisited: how frequent are attention shifts to one's name in an irrelevant auditory channel? Journal of Experimental Psychology: Learning, Memory, and Cognition, 21, 255.

Woods, K. J. & Mcdermott, J. H. 2015. Attentive tracking of sound sources. Current Biology, 25, 2238–2246.

Yin, P., Mishkin, M., Sutter, M. & Fritz, J. B. 2008. Early stages of melody processing: stimulus-sequence and task-dependent neuronal activity in monkey auditory cortical fields A1 and R. Journal of neurophysiology, 100, 3009–3029.

Zhang, W. & Ding, N. 2017. Time-domain analysis of neural tracking of hierarchical linguistic structures. NeuroImage, 146, 333–340.

Zhou, X., De Villers-Sidani, É., Panizzutti, R. & Merzenich, M. M. 2010. Successive-signal biasing for a learned sound sequence. Proceedings of the National Academy of Sciences, 107, 14839–14844.

